# Diffeomorphic Independent Contrasts for Ancestral Reconstruction of Shapes

**DOI:** 10.1101/2025.02.26.638985

**Authors:** Michael Lind Severinsen, Morten Akhøj, Rasmus Nielsen, Stefan Sommer, Christy Anna Hipsley

## Abstract

Ancestral reconstruction is a fundamental challenge in evolutionary biology, requiring methods that can capture complex morphological changes while accounting for phylogenetic relationships. Current approaches are based on linear assumptions that often oversimplify the spatial relationships between anatomical features and fail to account for landmark correlations within shapes. Here, we introduce a novel method that combines the ability of Large Deformation Diffeomorphic Metric Mapping (LDDMM) to model smooth, invertible transformations between shapes while preserving the relationships between landmarks with Felsenstein’s Independent Contrasts (IC) to iteratively reconstruct ancestral shapes along the branches of a phylogenetic tree. We call this method Diffeomorphic Independent Contrasts for Ancestral Reconstruction of Shapes (DICAROS). We validate DICAROS against two existing methods: (1) Linear predictors using Ordinary Least Squares and (2) Ancestral character estimation using maximum likelihood under Brownian Motion and apply DICAROS to a dataset of swallowtail butterfly species (Family Papilonidae, Order Lepiodetra) to reconstruct the ancestral shape and visualize evolutionary trajectories in a phylomorphospace from the contrasts. We conclude that DICAROS outperforms the existing methods in terms of accuracy and provides a more accurate reconstruction of the ancestral shape for non-symmetric phylogenetic trees. With DICAROS we show a transition between un-tailed and tailed papilinodae species while also illustrating how images of modern species would look under the DICAROS ancestral reconstruction

---

Evolutionary studies of morphology and omics have experienced rapid advancement in recent years, driven by new analytical methodologies and technological progress. From the increased accessibility of phylogenetic analyses and geometric morphometric (GM) through open-source software packages like Geomorph (Adams (2013, 2024)) and PhyTools (Revell (2024)) in the programming language R. These tools enable researchers to do phylogenetic comparative analysis while accounting for shared inheritance between related species in morphological and phylogenetic analyses, building on foundational work by Felsenstein and Bookstein (Felsenstein (1985); Bookstein (1997)).

Parallel advances in genomic sequencing and assembly (Höhna (2016); Zhang (2025)) have enabled phylogenetic inference across large evolutionary trees (Misof (2014); Kawahara (2023); Stiller (2024)). Which allows for the study of complex shape change in across tree of life, such as the mammalian skull evolution (Goswami (2023)) to bird brain and skull development (Chiappe (2024)) and butterfly wing vein patterns (Chazot (2016); Owens (2020)).

The increasing scale and complexity of these taxonomic datasets have been supported by the digitization of museum collections (Nelson (2019)), with resources like GBIF (GBIF (2024)), Morphosource (Morposource (2024)), and community-driven species mapping platforms (Inaturalist (2024)) providing unprecedented access to information across institutions. With these resources it is possible to combine phylogenetic information with shape data derived from landmarks placed on anatomical correspondences in 2D images or 3D reconstructions (Gunz (2013); Bardua (2019); Mitterocker (2021)). This enables ancestral shape reconstruction (Schluter (1997); Revell (2024)) and visualization in lower-dimensional phylomorphospaces (Polly (2013); Baken (2021)).

The current paradigm within morphological methods for ancestral reconstruction, including methods such as Ordinary Least Squares (OLS) linear predictors (Adams (2013, 2024)) and Ancestral character estimation using maximum likelihood under Brownian Motion (Revell (2024)), rely on linear assumptions that simplifies the complex spatial relationships between anatomical features. A key limitation is their inability to account for the correlation between landmarks within shapes, neglecting how changes in one landmark’s position can influence others during, e.g., reconstruction. This correlation structure is critical for two main reasons. First, biological shapes exhibit inherent modularity through physical proximity and functional relationships between features (Mitterocker (2007); Klingenberg (2008); Adams (2016)). Ignoring these relationships leads to oversimplified models that may miss important biological patterns. Second, understanding landmark correlations provides insights into developmental constraints and evolutionary mechanisms shaping morphological variation (Hallgŕımsson (2009); Young (2010)).

To address these limitations, we employ Large Deformation Diffeomorphic Metric Mapping (LDDMM) (Beg (2005); Younes (2010)). In this sense, our work can be seen as a continuation of Tangent Phylogenetic PCA (Akhøj (2023)), which is a method for ancestral state reconstruction based on nodes taking values on any finite-dimensional Riemannian manifold, including the LDDMM landmark manifold. LDDMM provides a mathematically rigorous framework for analyzing landmark configurations by explicitly modeling the correlations between landmarks. Operating as a Hamiltonian framework, LDDMM enables shape registration through smooth, invertible mappings that follow optimal geodesic paths between configurations. This approach preserves the topological relationships between landmarks throughout the transformation process, making it particularly suitable for biological shape analysis. The framework has been successfully applied in computational anatomy and medical imaging for tasks such as brain mapping (Durrleman (2014); Miller (2015)).

This study leverages LDDMM’s geodesic paths to reconstruct ancestral shapes along branches in a phylogenetic tree by combining LDDMM with Felsenstein’s *Independent Contrasts* (IC) method (Felsenstein (1985)). We call this novel approach *Diffeomorphic Independent Contrasts for Ancestral Reconstruction of Shapes* (DICAROS). Unlike traditional geometric morphometrics based on Kendall’s shape space (Klingenberg (2020)) and Procrustes alignment (Gower (1975)), the LDDMM framework offers several key advantages: it explicitly models landmark correlations, enables stochastic shape development models, and extends beyond landmarks to continuous curves, surfaces, and images. Our primary contribution is reformulating Felsenstein’s method using Riemannian manifold operations while maintaining equivalence to the original Euclidean approach when applied to vector data. We validate DICAROS by comparing the root reconstruction against two established methods: (1) Ordinary Least Squares (OLS) linear predictors (Adams (2013, 2024)) and (2) Ancestral character estimation using likelihood under Brownian Motion (Revell (2024)). To demonstrate DICAROS’ practical utility, we apply it to reconstruct ancestral wing morphology on swallowtail butterflies (Family Papilionidae, Order Lepidoptera), by showing the phylomorphospace estimated by DICAROS, and the evolutionary shape trajectories from a leaf image to the root shape.

## Materials and Methods

### Independent contrasts for LDDMM shape observations

The *independent contrasts* (IC) method (Felsenstein (1985)) reconstructs ancestral states and estimates evolutionary covariance matrices. Here, we generalize the IC to handle shape data, where each node value *x* represents a shape defined by landmarks *x*^1^, …, *x*^*k*^ ∈ ℝ^*d*^. Rather than treating shapes as vectors in Euclidean space ℝ^*d·k*^, we view them as elements of the LDDMM landmark manifold where distances between shapes are measured by geodesics paths. We combine LDDMM with IC by interpreting the phylogenetic tree as a network of geodesic paths on this manifold. Starting from the leaves, we traverse the tree in post-order, computing the diffeomorphism between sister taxa at each internal node. This diffeomorphism defines a geodesic path between the branches of shape *x_i_* and *x_j_*. Given the relative position of their common ancestor *y_i_* along this path, we reconstruct the ancestral shape by deforming *x_i_* toward *x_j_* and stopping at the appropriate position of *y_i_*. Where *x_i_* corresponds to the child of *y_i_*, with the shortest branch lenght. We repeat this process until reaching the root.

The LDDMM framework enables the computation of an evolutionary covariance matrix through the right Lie transport of cotangent vectors, accounting for the nonlinear geometry of shape space. This allows us to perform a generalized version of Phylogenetic PCA (Revell (2009)) that respects the manifold structure of the shape data. The key advantage of our approach is that both the ancestral reconstruction and covariance estimation are performed using operations that are appropriate for shape data, unlike traditional methods that rely on linear approximations. The LDDMM framework introduces a kernel parameter *σ* which determines the spatial scale of correlations between landmarks through a Gaussian kernel *κ*(*q_i_, q*_*j*_) := exp^−∥*q*^*^i^*^−*q*^*^j^* ^∥2/2σ2^ ∈ ℝ with parameter *σ >* 0 (Miller (2002); Younes (2010); Pennec (2019)). A larger kernel width means that movements of one landmark have a broader influence on surrounding landmarks, while a smaller width allows more localized, independent deformations. This parameter effectively controls the “stiffness” or “elasticity” of the deformation field but also implies there is no canonical metric on the LDDMM landmark manifold, making *σ* a hyperparameter.

The LDDMM landmark manifold is a Riemannian manifold, where standard vector space operations like addition and multiplication are not available. Instead, the *Riemannian exponential* map Exp_x_(*v*) and *Riemannian logarithmic* map is used Log_x_(*y*) as generalizations of addition and subtraction. The logarithm Log_*x*_*_i_* (*x_j_*) between two shapes *x_i_* and *x_j_* yields a vector *v* in the *tangent space T_x__i_* *R*^*d*×*k*^ at *x_i_*, providing a local linear approximation of the curved manifold (Michor (2020)). The exponential map Exp_*x*_*_i_* (*x_j_*) takes this tangent vector and maps it back to a point on the manifold.

When computing these maps for independent contrasts, the kernel width *σ* directly influences how shape differences are interpreted through the cometric 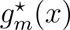, which is a block matrix of Gaussian kernels, here we choose *σ* as the mean distance between each landmark and its nearest neighbor. Since the contrasts are computed as cotangent (momentum) vectors in different cotangent spaces 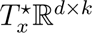, we need to map them to a common space to compute the phylogenetic covariance matrix. For the LDDMM landmark manifold, this mapping between cotangent spaces can be achieved by composing linear transformations based on the Jacobians of the optimal diffeomorphisms along the geodesic path between shapes (see Supplementary Section 1 for mathematical details).

To adapt the IC method for this manifold structure, we modify the computation of covariance matrices by replacing standard vector outer products with Riemannian metric-weighted outer products, as shown in Equation (1). Our algorithm for IC on the LDDMM manifold follows the same structure as the Euclidean version, differing only in three key steps that employ Riemannian operations: step *3* (raw contrast computation between nodes), step *5* (weighted mean computation for inner nodes), and step *9* (evolutionary covariance matrix computation using standardized contrasts) as detailed in Algorithm 1. With the evolutionary covariance matrix we project all the evolutionary shape trajectories with Principal Component Analysis (PCA) into a so-called Phylomorphospace.

#### Algorithm 1 Independent contrasts on the LDDMM landmark manifold

**Require:** A phylogenetic tree with |*V* | nodes, |*E*| branches and leaf node observations

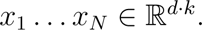

**Ensure:** Estimates of the root value and evolutionary covariance matrix.

1. **while** |*V* | *>* 1 **do**
2. Choose two leaf nodes *x_i_* and *x_j_* with common parent node.
3. Compute the raw contrast *c_ij_* = Log_x*i*_ (*x_j_*), where Log is the Riemannian logarithm.
4. Compute the standardized contrast

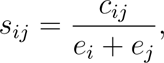

where *e_i_, e_j_* are the lengths of the branches leading to their common parent node.
5. Remove nodes *i, j* and branches *e_i_, e_j_* from the tree, so that |*V* | is lowered by 2. The parent node *k* now becomes a leaf. Assign to it the node value

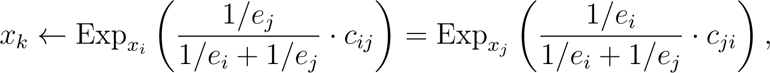 where Exp is the Riemannian exponential.
6. Increase the length of the edge leading from node *k* to its parent as follows,

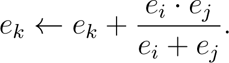
7. end while
8. Let 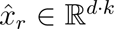 be the assigned node value of the single remaining node in the tree. This is the root estimate.
9. Let *s*_1_, …, *s*_|_*_E_*_|/2_ be the standardized contrasts computed above. Map each contrast to the tangent space at 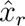 and denote the resulting vectors by 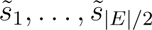 (see Supplementary Section 2 for details on this map). The estimated evolutionary covariance matrix is

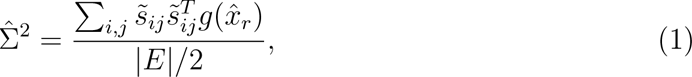

where 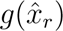 is the Riemannian metric matrix at 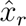.

**return** 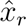 and 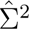

Given two landmark shapes *x_i_* and *x_j_*, the geodesic contrast *c_ij_* = Log_x_*_i_* (*x_j_*) defines a geodesic path of shapes from *x_i_* to *x_j_*. This geodesic path can be extended beyond the landmarks to include all points in the surrounding domain, such as a rectangle encompassing both shapes. The entire domain is transformed by a diffeomorphism *ϕ* that maps the *k* landmarks of shape *x_i_* to their corresponding landmarks in shape *x_j_*, i.e., 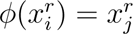 for *r* = 1, …, *k*. The movement of surrounding points is determined smoothly _i j_ through an interpolation kernel (see Supplementary Section 2). In practice, the Gaussian kernel width parameter *σ_DICAROS_*.

For image reconstruction along the phylogenetic tree, consider *x_i_* as the estimated parent node of a child node *x_j_*. If *x_j_* has an associated image of *G* pixels {*p_i_*}_i=1,…,G_ arranged on a rectangular grid around its landmarks, we construct the parent node image as follows: For each pixel *p*^′^_i_ in the parent image, we compute its color as a weighted average of the colors of nearby pixels in the child image after applying the transformation *ϕ*(*p*^′^_i_). This process propagates iteratively along the branches of the tree, ultimately yielding the reconstructed image at the estimated root shape and visualizing the evolutionary trajectory through the tree.

In terms of implementations, DICAROS builds upon the numerical differential geometry algorithms contained in JaxGeometry (Kühnel (2017, 2019)). These algorithms are implemented using JAX (Bradbury (2018)), a library that enables highly optimized and efficient numerical computations. The phylogenetic components of our method, including the crucial “Upwards Pass” algorithm, which transverse the tree from leaves to root, are implemented in Hyperiax https://github.com/ComputationalEvolutionaryMorphometry/hyperiax. Hyperiax leverages JAX to efficiently process independent branches in the phylogenetic tree in parallel, significantly boosting computational performance.

### Shape simulation and comparison of method

To evaluate DICAROS, we simulate shape evolution along phylogenetic trees using the Kunita Flow(Stroustrup (2025)). The simulated leaf shapes are then used to reconstruct the root shape, allowing us to compare DICAROS reconstruction ability, and furhter compare against established reconstruction methods. The Kunita flow provides key advantages over a Brownian Motion (BM) model, which would likely produce biologically unrealistic shapes(Diaz-Uriarte (1996)). Instead, the Kunita flow models a shape as a set of *k* landmarks *x*_1_*, …, x_k_* in a domain *D* ⊂ ℝ^d^ (for *d* = 2 or 3), where landmark movements are spatially correlated based on their proximity. This correlation ensures that nearby landmarks move coherently and that shapes maintain biological plausibility as they evolve (Arnaudon (2019, 2021); Sommer (2021); Stroustrup (2025)). The spatial correlation is modeled using a scaled Gaussian kernel 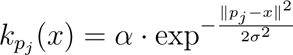, where *σ* controls the spatial extent of correlation (kernel width) and *α* determines the size of the process.

We benchmark DICAROS against two standard approaches in the field: (1) Ordinary least squares estimation using *gm.prcomp* and *shape.predictor* from Geomorph v4.0.6 (Adams (2013)) and (2) Maximum likelihood estimation of ancestral states under Brownian motion using *anc.ML* from Phytools v2.3-0 (Revell (2024); Schluter (1997)). For fair comparison, all reconstructed shapes are aligned to the original root via general Procrustes alignment (Rohlf (1990)) using *gpagen* from Geomorph (Adams (2013, 2024)).To evaluate reconstruction accuracy, we compare the reconstructed root shapes to the original root using normalized Procrustes distance. Specifically, we calculate the mean least squared error (MLSE) between corresponding landmarks using Equation 2:

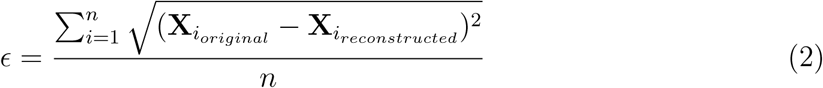

where *ɛ* is the error, **X**_i_ is the coordinate vector of landmark *i*, and *n* is the total number of landmarks. The relative performance between methods is quantified as *^ε^* ^= *ɛ*^**_establish method_** ^− *ɛ*^**_DICAROS_**.

### Biological data

As a biological application of DICAROS, we analyzed Lepidoptera wings from the Papilionidae family. The phylogenetic tree was derived from (Kawahara (2023)) and matched with image data through the Global Biodiversity Information Facility (GBIF.org) (GBIF (2024)). To ensure data quality, we restricted our selection to museum collections of male specimens to account for sexual dimorphism. Images showing damaged wings or improper orientation were manually excluded.

The phylogenetic tree was pruned using the *drop.tip* function from Ape (Paradis (2019)) to remove taxa without shape data. We used *Segment Anything* (SAM) (Kirillov (2023)) with *Grounding Dino* (Liu (2023)) to automatically extract the butterfly from its background, using the prompt “Butterfly”. Images were cropped to include only the butterfly. Cases where SAM failed to identify a butterfly or identified multiple specimens were excluded.

To differentiate between morphological features, hind-/fore-wings and thorax, six anatomical landmarks were annotated on the outline for each image in a similar setting as seen in (Chan (2022)). The annotation was done with the software *Supervisely*, by external consultants hired through *fiverr.com* The butterfly contour was extracted using *cv2.findContours* from the Python package *opencv* (Bradski (2008)). The contour was matched with the six anatomical landmarks, and for the regions covering the wing sections, 30 equidistance landmarks were equidistance resampled with interpolation on the wing contour The methodology is detailed in Supplementary Section 3.

Each specimen was manually categorized based on tail morphology into three categories: untailed, tailed, or long-tailed. Using DICAROS, a full ancestral reconstruction was executed for the entire phylogeny, and the morphological changes from the leaves to the root. Further, we use the covariance to visualize the entire evolutionary trajectory by projecting the covariances and shape into the phylomorphospace.

## Results

The evaluation of DICAROS is based on the simulated leave shapes from applying Kunita Flow on different phylogenies with different parameters, root shapes, and sizes. Figure 1 shows the steps of this comparison. First, a root shape (here, a butterfly) is simulated on a phylogeny with four leaves. The leaf shapes are then used to reconstruct the root shape and finally compare the reconsturcted root the original root shape.

**Fig. 1:**
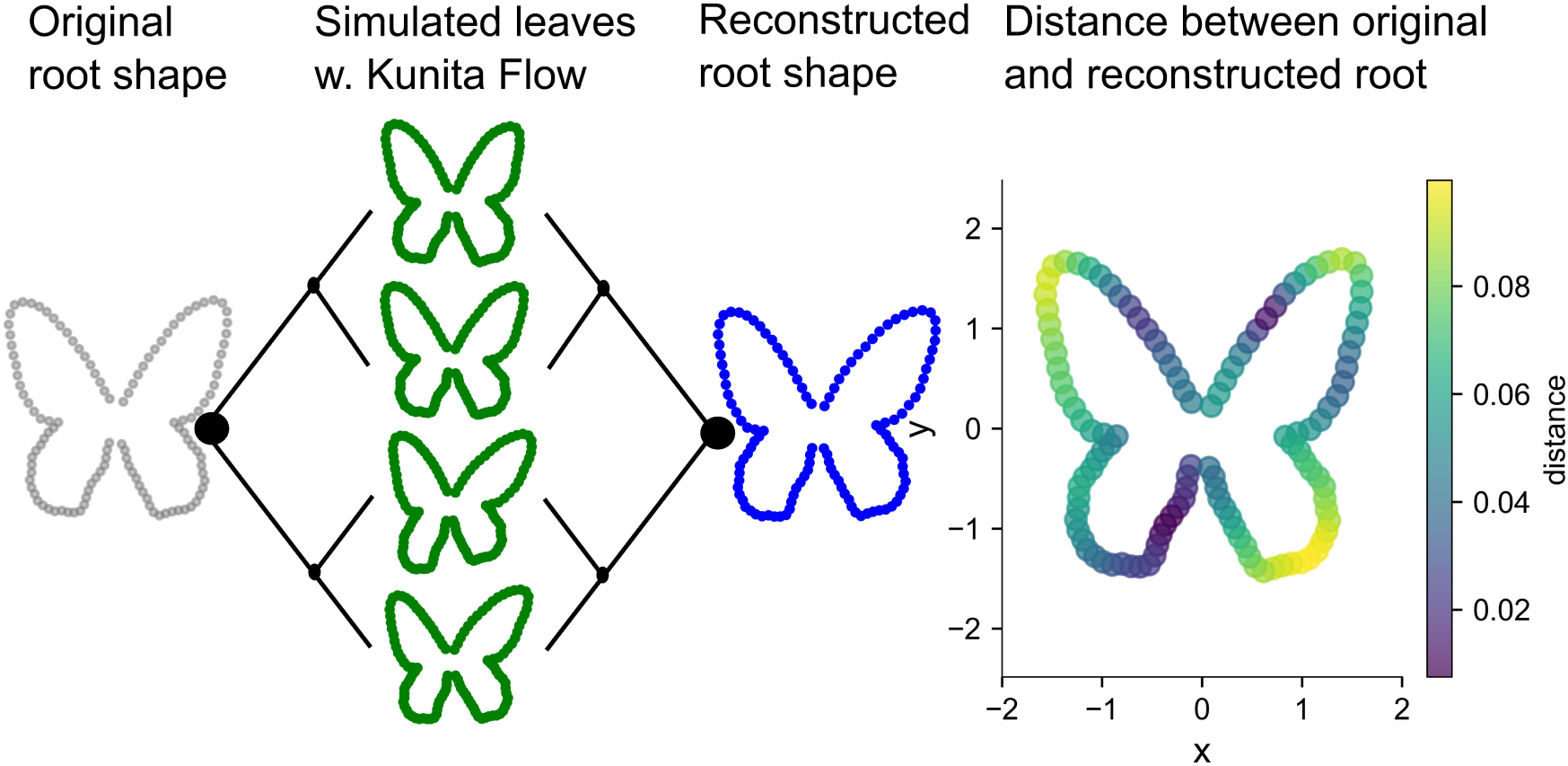
The grey butterfly is used as the root shape for the Kunita Flow on a symmetric ultrametric phylogeny with four leaves. The root shape is simulated with Kunita Flow on the phylogeny to obtain the green leaf shapes. These leaf shapes are used for the reconstruction of the root shape using DICAROS on the same phylogeny. The reconstructed root shape in blue is then compared to the original root shape using the Euclidean distance between each landmark in the reconstructed and original root shape

To repeat this experiment, four different root shapes are used. Two calculated shapes (a circle and a sphere) and two real shapes (a butterfly and a bird beak), described in Table 1.

**Table 1:** The four shapes and their dimension (d) and number of landmarks (m), and the origin of the root shape

| Shape | (d,m) | origin |
| --- | --- | --- |
| Circle | (2,30) | Calculated |
| Sphere | (3,50) | Calculated |
| Butterfly | (2,118) | *Atrophaneura dixonii* (own data) |
| Birdbeak | (3,79) | *Fratercula arctica* (Cooney (2017)) |

The robustness of DICAROS is tested by varying the *σ_DICAROS_* by a factor of 2 and 10, with *σ_DICAROS_* being the mean squared distance to the closest neighboring landmark. The test is executed on both symmetric and asymmetric ultrametric trees with a fixed total length. When the tree size is increased, the individual branch length decreases to represent a greater species diversification within the same period of time. The tree size vary by2^k^, (*k* = 2, .., 8) leaves and is repeated for all four shapes. For each configuration, the simulation is repeated 100 times. (Fig. 2)

**Fig. 2:**
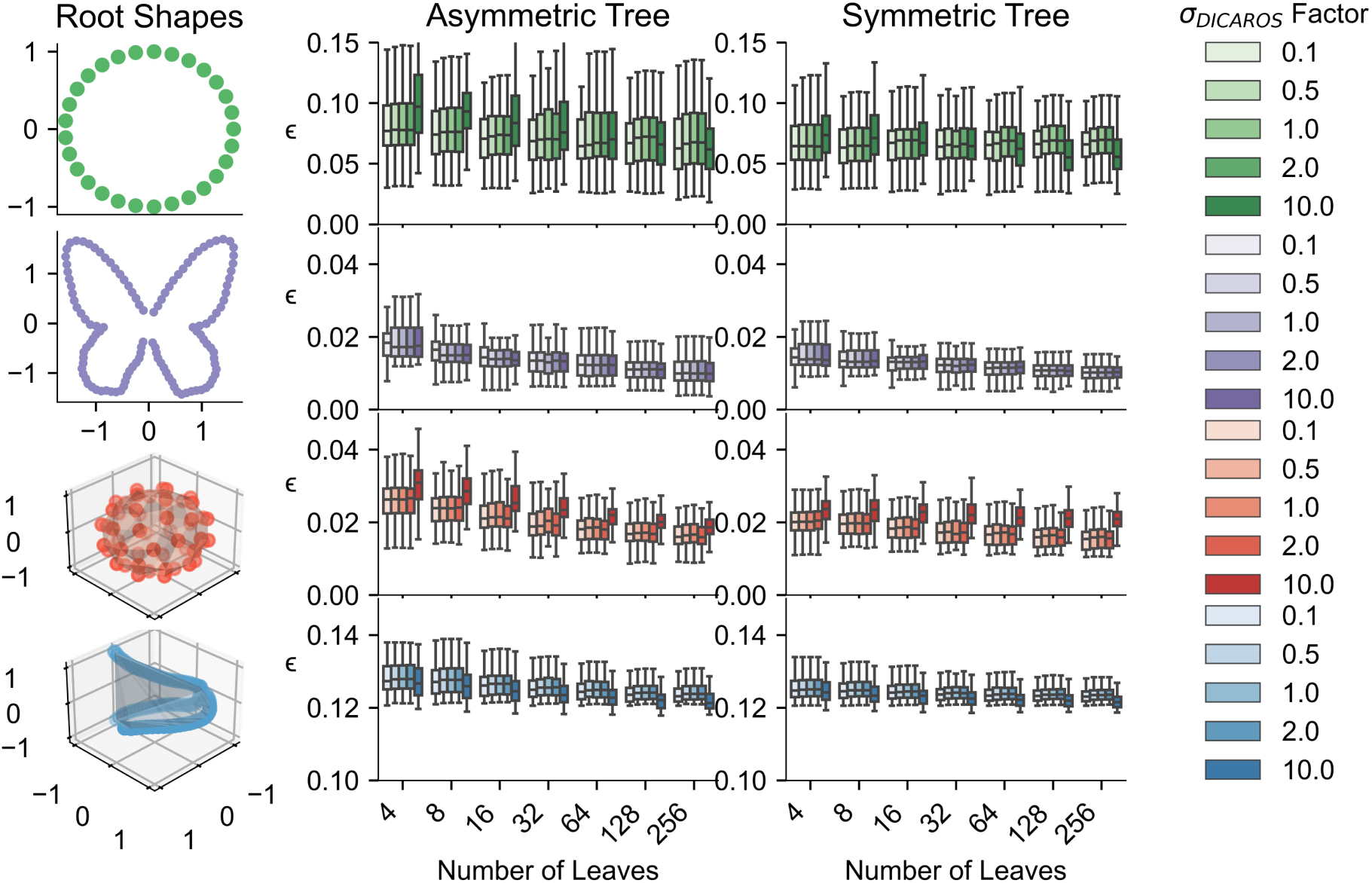
The landmark configuration of each root shape is shown to the left, with the MSLE between the original and reconstructed root for both types of phylogenies, tree sizes, and scaled *σ_DICAROS_*. The Y-axis is adjusted for each root shape, but the tendencies show a higher variance for the MSLE under an asymmetric phylogeny compared to the symmetric phylogeny

Figure 2 shows the robustness of the method for different *σ_DICAROS_*, and the MSLE remains consistent between symmetric and asymmetric phylogeny across all 100 repetitions. We denote *σ_DICAROS_* as not to confuse it with the kernel width in the Kunita Flow, noted as *σ_KF_*. Figure 2 demonstrates the robustness of the method regardless of tree topology. Based on these results, we fixed sigma at its base value without applying scaling factors for subsequent analyses.

Continuing, with a fixed *σ_DICAROS_* at the factor equal to one, the experiment is repeated for different parameters of the Kunita Flow. Here, the variance and noise parameter (*σ_KF_*, *α*) is changed to illustrate less and larger shape changes across the phylogeny. For each simulation, the root is now reconstructed using DICAROS, OLS, and BM. For each evaluation, the relative procrustes distance *ɛ* = *ɛ***_establish method_** − *ɛ***_DICAROS_**, is shown. Where a positive value on the y-axis, indicates a better reconstruction of the root shape by DICAROS, compared to the other method. The relative error between OLS and DICAROS is shown in Figure 3, where DICAROS are more accurate than OLS for asymmetric chronograms, but they are similar for the symmetric chronogram. Except a few outliers for the sphere where the OLS is not able to reconstruct the root as good as for the other simulation.

**Fig. 3:**
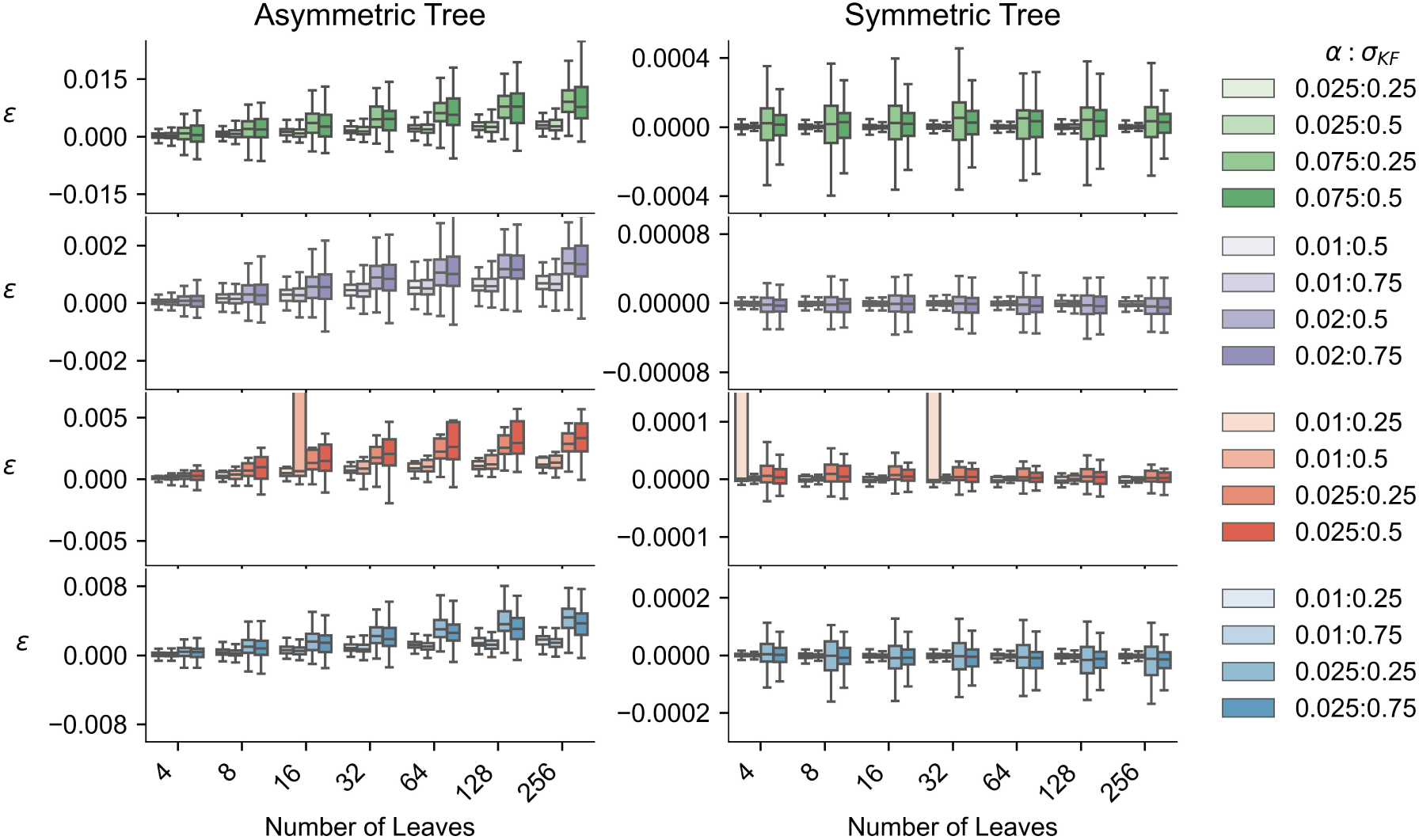
The relative distance *ε* = *ɛ***_OLS_** − *ɛ***_DICAROS_** is shown for four different root shapes (circle, butterfly, sphere, and bird beak) under varying Kunita Flow parameters. Positive values indicate that DICAROS outperforms OLS, which is seen for the asymmetric trees and increasing the tree sizes. For symmetric phylogenies (right column), both methods perform similarly with error distributions centered around zero. The boxplots show the distribution of relative errors across 100 simulations for each configuration

The relative error between methods shown in Figures 3 and 4 corresponds to ≈ 10% of the total error magnitude, when comparing the base case (*σ_DICAROS_* factor = 1) in Figure 2, which shows absolute MSLE values for the circle is within [0.05;0.15], to the relative differences in Figures 3 and 4 within [-0.005;0.015]. We observe this consistent 10% relationship for all root shapes. This indicates that while DICAROS shows improvement over existing methods, the magnitude of improvement is moderate compared to the overall reconstruction error.

**Fig. 4:**
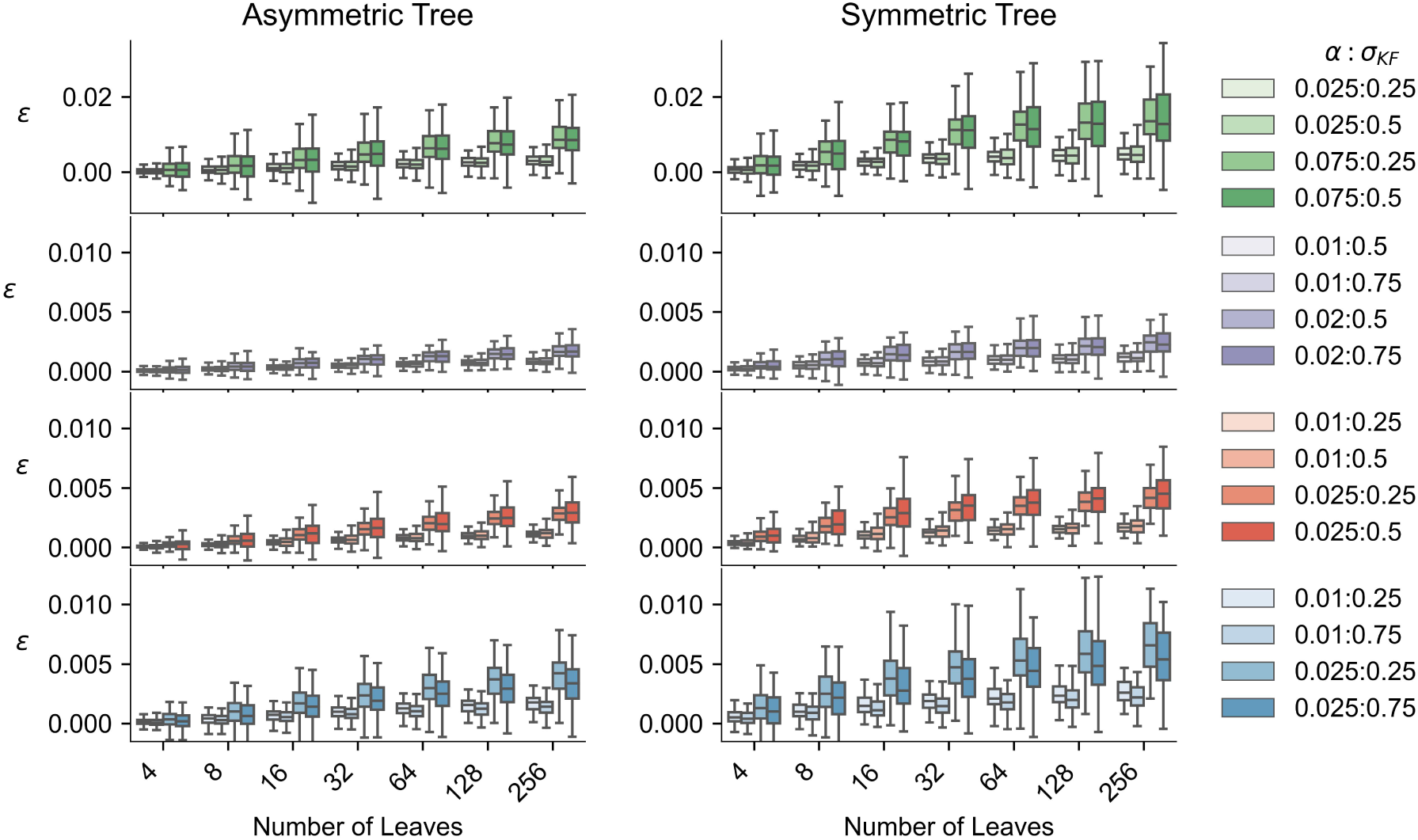
Comparison of reconstruction accuracy between the Ancestral character estimation using maximum likelihood under Brownian Motion and DICAROS using relative MSLE (*ε* = *ɛ***_BM_** − *ɛ***_DICAROS_**). The consistently positive values across both symmetric and asymmetric phylogeny indicate that DICAROS achieves better reconstruction accuracy compared to this method. This pattern holds true across all tested shapes and parameter configurations, demonstrating the robust performance advantage of DICAROS

### Diffeomorphic Independent Contrasts for wing shape analysis

The butterfly phylogeny with example images and shapes is shown in Figure 5. The entire dataset encompasses 992 individual butterflies distributed across the 49 Papilionidae species. The phylogenetic analysis contains three main morphological categories based on hindwing tail length: untailed, tailed, and long-tailed species. All mean shapes can be found in Supplementary Section 5 and origins of all images can be found in Supplementary Section 6. The entire dataset is shared in DataDryad; https://doi.org/10.5061/dryad.41ns1rnr7

**Fig. 5:**
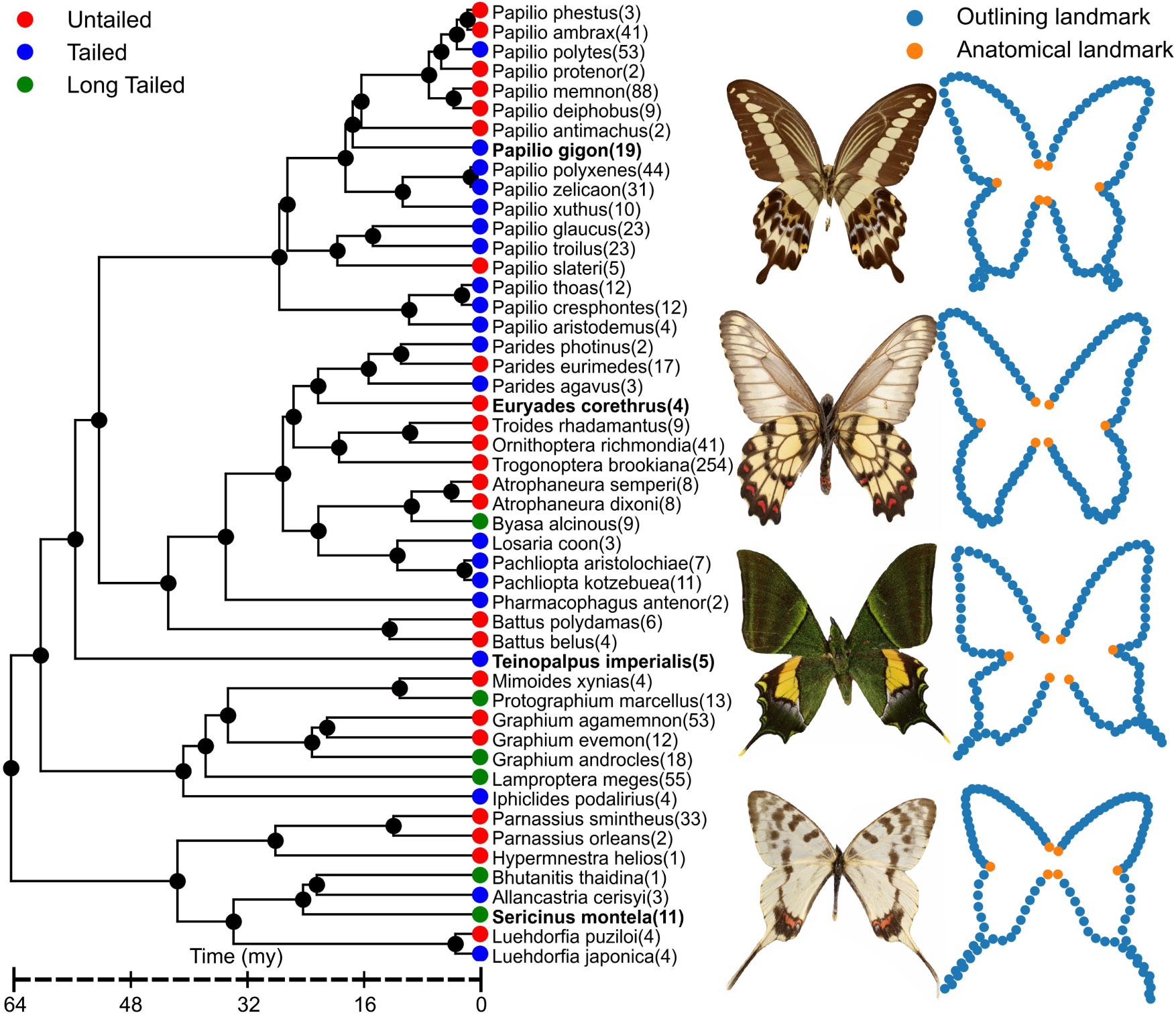
Phylogenetic tree of the 49 Papilionidae species in our study, adapted from the larger Lepidoptera phylogeny from (Kawahara (2023)), with the number of specimens pr species in parenthesis. Species are categorized based on hindwing tail length: untailed, tailed, and long-tailed, with examples of four species specimens marked in bold *Papilio gigon*, *Euryades corethrus*, *Teinopalpus imperialis* and *Sericinus montela*.

With the mean shape of each species, we can visualize the phylomorphospace of the butterfly wing shape. The phylomorphospace is shown in Figure 6, where it illustrates the transition from the untailed to the long-tailed species. The color of the leaves matches the color of the species in Figure 5, and the root is placed as a star. For each branch in the phylogeny, a grey line is drawn and selected leaf mean shapes is shown across the path of least resistance.

**Fig. 6:**
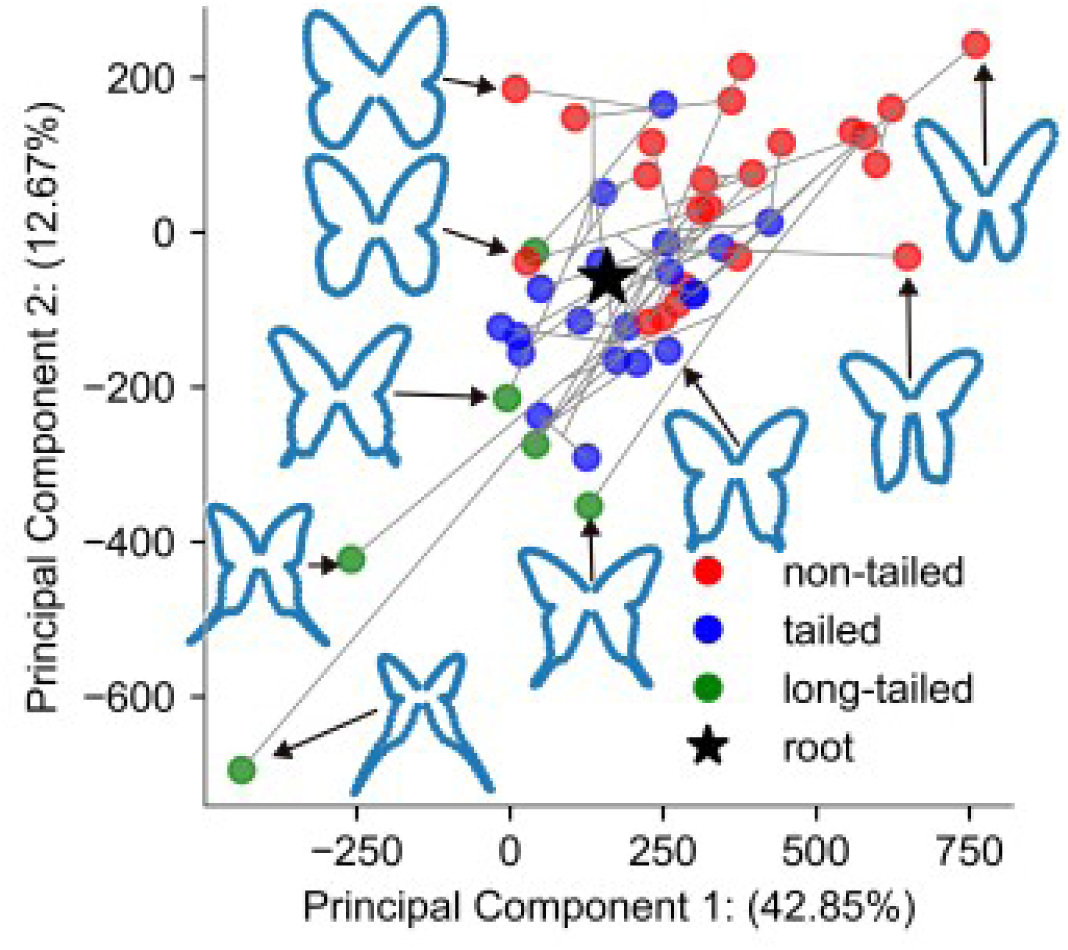
The DICAROS Phylomorphospace of butterfly shapes across the Papilionidae phylogeny. The plot shows the distribution of species in the morphological space, with the ancestral root state marked as a star. Grey lines represent phylogenetic branches connecting species, and colors correspond to tail morphology categories as defined in Figure 5. The visualization reveals the transition of shape between long-tailed and untailed species

DICAROS reveals distinct evolutionary patterns in butterfly wing morphology demonstrating a clear separation between long-tailed and untailed species. To visualize the evolutionary transitions, we applied the estimated deformation fields from DICAROS to transform modern specimen images along their phylogenetic branches back to the root. Figure 7 shows this process for eight individuals across different species. The full animation, going from the leaf shape and continuous to the root shape, is visible at the GitHub supplementary repository https://github.com/MichaelSev/DICAROS.

**Fig. 7:**
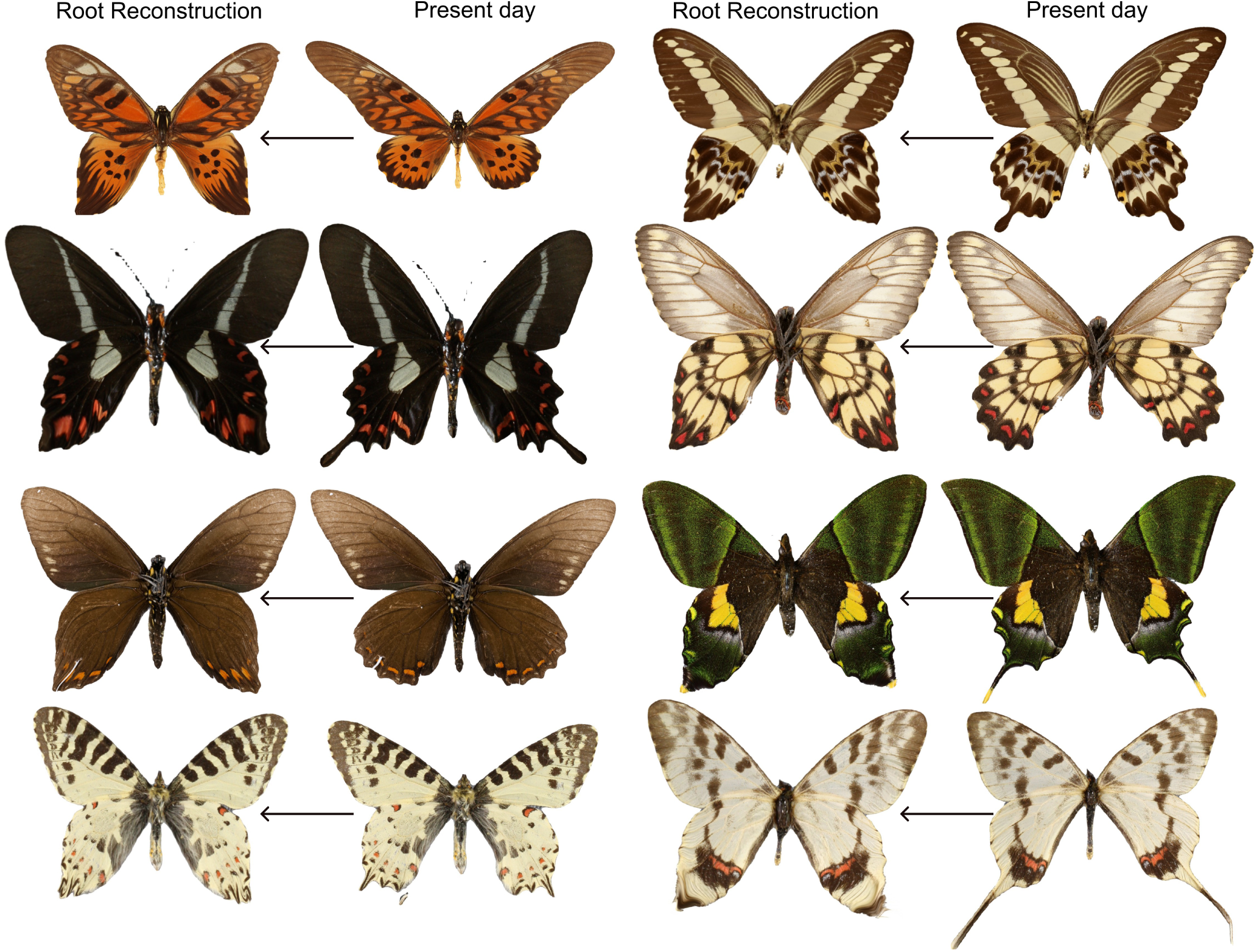
The application of DICAROS on the image for the eight individuals across different species. With the right column corresponding to the individuals from Figure 5, *Papilio gigon*, *Euryades corethrus*, *Teinopalpus imperialis* and *Sericinus montela* and the right column corresponding to these species in bold; *Papilio antimachus*, *Parides agavus*, *Battus belus* and *Allancastria cerisyi*. The origin of the species is detailed in supplementary materials. A full transformation for the leaf species to the root species can be found at https://github.com/MichaelSev/DICAROS

## Discussion

DICAROS represents an advancement in the field of ancestral shape reconstruction by demonstrating the novel use of LDDMM in a phylogenetic setting, utilizing tree topology as a map to reconstruct ancestral shapes. A key feature of DICAROS is its ability to incorporate correlations between landmarks, allowing for reconstruction to be applied to entire images through image manipulation (Fig. 7). Additionally, if the landmark data are only aligned using translation and rotation, the ancestral size can also be reconstructed.

Using the methodology from Independent Contrasts(Felsenstein (1985)), it estimates the evolutionary covariance matrices to project shape data of both leaves and reconstructed inner node/root into phylomorphospace. The DICAROS implementation utilizes modern frameworks including *JAX* (Bradbury (2018)), *Jaxgeometry* (Kühnel (2017, 2019)), and *Hyperiax*, providing good performance and stability across different dataset sizes and shape complexities. The application of DICAROS to butterfly wings, combined with semi-automated segmentation and landmarking tools (Kirillov (2023)), not only provided insight into the tail evolution/devolution of the swallowtail but also demonstrates the potential for creating larger biological geometric morphometric datasets with greater time efficacy compared to traditional human labor approaches.

DICAROS naturally operates under a design constraint that can only be used with a rooted phylogeny. In contrast, other methods can illustrate the morphospace with and without phylogeny(Baken (2021)). Furthermore, the implementation of DICAROS only uses bifurcating nodes, but the method can be generalized to include multifurcating nodes. For the bifurcation nodes, the leaves can also be represented from a fossil or prehistoric species, these tips can be inserted in the already existing inner nodes to make it the socalled total evidence phylogeny. This will likely make the inner nodes more exact in their reconstruction. Using the iteration of local computations that form the IC algorithm it is a computational advantage instead of simply evaluating the closed-form MLE estimates. When dealing with large phylogenies, DICAROS avoids the inversion of large matrices, as seen for the MLE estimates. In the case of manifold valued nodes, as assumed by DICAROS, the iteration of local computations is an advantage since the local computation of inner nodes (step *5* in Algorithm 1) can be solved by a single evaluation of the exponential map without the need of larger optimization schemes that would otherwise be necessary for estimating the root directly from the leaves. The latter is done in Tangent Phylogenetic PCA Akhøj (2023)), which formulates the root estimate as a weighted Frechét mean (i.e. a mean on a Riemannian manifold), the computation which demands the the same number of exponential map evaluations as the number of leaf nodes at every iteration until convergence. This number of exponantial map evaluations well exceed that of DICAROS.

To evaluate the performance of DICAROS relative to existing methods, we conducted four simulation experiments (Fig. 1. First, we simulated shapes evolving along phylogenetic trees using the Kunita flow. Secondly, these simulated leaf shapes were used to reconstruct the root shape using three methods: DICAROS, Ordinary least squares(Adams (2013, 2024)), and Brownian Motion Maximum Likelihood(Revell (2024)). Third, we compared the reconstructed root shapes to the true ancestral shape to calculate reconstruction error (Eq. 2), and finally repeated this process under different topologies, tree sizes, and root shapes to assess method performance across varying conditions (Fig 3,4). We show that DICAROS perform slightly better in ancestral shape reconstruction under an asymmetric chronogram and for increasing tree sizes (Fig. 3,4). The results were close to identical for a symmetric chronogram (Fig. 3 relative to the OLS from Geomorph and DICAROS are more accurate in shape compared to the BM(Fig 4). This may be related to the reconstruction under a symmetric tree roughly corresponds to the equal weighted average among species. Whereas for the asymmetric tree, the morphological evolution and tree topology is crucial, because its determine how the evidence from different branches are weighted in the reconstruction process.

While our simulations provide insights into comparing DICAROS with existing methods by the ancestral root shape, it is not possible to directly evaluate evolutionary covariance estimation, as the Kunita flow used does not have a parameter directly corresponding to the estimated evolutionary covariance matrix (Equation 1). Furthermore The Kunita Flow arrives from a stochastic process tailored for shapes rather than a BM model. When we evaluate our method based on simulations, we note that neither the standard IC nor the DICAROS estimates correspond to maximum likelihood estimates under this stochastic model, and therefore, is it fair to compare DICAROS and the other methods with these simulated shapes.

To evaluate the sensitivity of DICAROS to the kernel size, we tested different values of *σ_DICAR_* by varying the default value (average distance of neighboring landmarks) by factors of 2 and 10. Note that this *σ_DICAR_* does not correspond to the kernel size used in the simulations (*σ_KF_*). The method showed robust performance across a range of kernel sizes (Fig. 2). With very small kernel sizes approximating Euclidean space, performance remained similar to the default value, demonstrating that the combination of LDDMM with phylogenetic shape estimation provides better ancestral reconstruction compared to OLS and BM models. However, performance degraded with excessively large kernel sizes, as these caused all landmarks to move uniformly rather than capturing local shape differences. This occurs because large kernels encompass most or all landmarks simultaneously, preventing the LDDMM method from modeling independent landmark movements needed for accurate shape reconstruction.

Butterflies are often used as model organisms in fields of biology, such as conservation biology (Collins (1985); Śanchez-Bayo (2019)), phylogenetic inference (Caterino (1999); Kawahara (2023)) and studies of independent evolution between fore- and hind-wings (Chai (1990); Chotard (2022)) and how to do high throughput imaging of museums specimens (Chan (2022)). Here we focus on the swallow-tailed family Papilonidae, which has long fascinated researchers due to their many speciation events and mimicry abilities (Mallet (1998); Carvalho (2024)), morphs (Le Roy (2019)), and sexual dimorphism (Condamine (2012)). The tails appear independently throughout the phylogeny and are not restrained to a monophyletic group. The tail can most often be on the fourth vein, but this also differs in placement, count, shape, and occurrence between species within and across genera and sexes (Nakae (2021)). Here, we aim to reconstruct the trajectory of shape evolution for the swallowtail family using DICAROS and illuminate that change using modern species images.

The biological dataset comprises wing shape data from 49 species of Papilionidae butterflies (Fig. 5), representing ≈ 20% of the recognized species in this family (Condamine (2023)), and ≈ 68% of the Papilionidae family from Kawahara et.al. 2023 (Kawahara (2023)). After sorting the obtained images through GBIF, we obtained a total of 992 specimens on to base our mean shape. While our sample size for some species is limited, it provides a foundation for exploring shape evolution within this group while showcasing the power of the DICAROS method. The application of semi-automated landmark placement on butterfly wing outlines utilized state-of-the-art segmentation tools (Kirillov (2023))(Fig. 5), together with using the contour instead of vein points with none/less homology across the family allowed us to showcase a more comprehensive representation of Papilionidae with a high number of individuals. For visualization purposes in the phylomorphospace, we categorized hindwing types into three groups: untailed, tailed, and long-tailed. While this classification simplifies the presentation in Figure 6, it is important to note that tail length exists on a continuous spectrum rather than as discrete categories. For this purpose the “tailed” group represents an intermediate state that shows overlap with both untailed and long-tailed morphologies. This categorization was used solely for visualization and was not incorporated into any quantitative analyses.

The phylomorphospace analysis (Fig. 6) revealed distinct patterns in the distribution of butterfly wing shapes across Papilionidae. It has a clear separation between long-tailed and untailed species, with a gradual transition of tail length in between. This indicates that the presence of tails is a factor in shape variation with small tail length representing intermediate evolutionary stages towards the long tail. With the root shape represented as tailed and is within the tailed group. This conforms with (Chotard (2022)), who employed a binary classification of tailed/untailed species with landmarks placed manually on each vein. The image reconstruction (Fig. 7) provides a visualization of shape evolution across evolutionary time. While this visual reconstruction effectively captures changes in wing morphology, it is important to note its limitations. The reconstruction is based solely on landmark shape data and does not account for other aspects of wing morphology, such as pigmentation patterns or wing-scale structures. The color scheme in the deformations is arbitrary and does not reflect actual evolutionary changes in wing coloration, but only from the leaf image. Despite these constraints, this approach represents a significant advancement in the quantitative analysis and visualization of morphological evolution in Lepidoptera. The reconstruction supports the hypothesis that the ancestral state of Papilionidae was likely tailed.

## Conclusion

In conclusion, this study introduces DICAROS as a novel method using LDDMM for ancestral shape reconstruction in phylogenetic contexts. The method offers significant advantages, including the incorporation of correlations between landmarks. While computationally more intensive than traditional approaches, DICAROS leverages modern tools like JAX, Jaxgeometry, and Hyperiax to enhance performance, mainly through GPU acceleration. As such, it represents an advancement in the field of geometric morphometrics evolution studies.

Our validation through simulations demonstrated that DICAROS obtained similar results and outperformed established methods in the case of asymmetric trees(Fig 3,4). The method’s accuracy improved with increasing numbers of leaf nodes, highlighting its potential for large-scale analyses. These simulations provide a solid foundation for the method’s reliability in reconstructing ancestral shapes while preserving both form and shape.

Our application of DICAROS to Papilionidae butterfly wing shapes supported insights into their evolution. We hypothesize that the evolution of modern non-tailed species originates from a tailed ancestor and that similar structures in phylomorphospace (Fig. 6) are observed for independence of hindwing and forewing shape evolution. The novel implementation of image reconstruction by DICAROS offers a new tool for studying any shape evolution on a phylogeny, which will contribute to our understanding of shape evolution across time.

## Acknowledgments

We express our gratitude to GBIF.org for its extensive work in assembling information across diverse organizations and data structures. We also acknowledge the valuable contributions of organizations, curators, taxonomists, and authors across museums who classified and photographed the butterflies. Our work has been enhanced by the tools developed by Google (JAX) and Meta (Segment Anything), which greatly improved our computational capabilities. We are also grateful to the authors of Geomorph for developing their R package and for their openness and fast responses. We also express our gratitude for software engineers and staff at AI companies, who build and designed; auto completions, spelling checkers and text generators for speeding up both coding and writing.

## Funding

This work was supported by a research grant (VIL40582) from VILLUM FONDEN and Danmarks Frie Forskningsfond (Independent Research Fund Denmark) grant ID 10.46540/3103-00296B.

## Data Availability

Code for figures and both simulated and biological dataset are shared in the data dryid DOI; https://doi.org/10.5061/dryad.41ns1rnr7. Further illustration and biological examples is available from Github repository; https://github.com/MichaelSev/DICAROS. For Hyperiax, we advertise for https://github.com/ComputationalEvolutionaryMorphometry/hyperiax, to follow the most recent updates.

## Notes

### Competing Interest Statement

The authors have declared no competing interest.

