## Supplementary Material for "Diffeomorphic Independent Contrasts for Ancestral Reconstruction of Shapes"

### 1 LDDMM and mapping between cotangent spaces

In this section we introduce the LDDMM Riemannian metric for landmark shapes. Parts of the text assumes a basic familiarity with Riemannian geometry. For an exposition of this in relation to statistical applications, see [1].

For our purposes, the *Large deformation diffeomorphic metric mapping* (LDDMM) framework can be seen as a way to define a Riemannian metric $g_m$ on $\mathbb{R}^{d\times k}$ that seems reasonable when a point in $\mathbb{R}^{d\times k}$ is interpreted as a collection of $k$ landmarks, i.e. a landmark matrix. A Riemannian metric determines a distance metric, which in the LDDMM case yields a sensible way to measure distances, and compute shortest paths, between shapes. What we refer to as the *LDDMM landmark manifold* is the Riemannian manifold $(\mathbb{R}^{d\times k}, g_m)$. We give more details on the metric below, but for now we mention that this manifold is geodesically complete, and so the induced geodesic distance metric $d_{g_m}$ on $\mathbb{R}^{d\times k}$ is complete (see [2] and [3]). Since this is a distance metric on $\mathbb{R}^{d\times k}$ it assigns non-zero distance between landmark matrices $x, x' \in \mathbb{R}^{k\times d}$ that represent the same shape, in the sense of Kendall's shape space, i.e. points $x, x'$ that are simply a rotation, scaling and/or translation of each other. Therefore a pre-processing step is necessary; a given set of observed landmark shapes are first Procrustes aligned before treating them as points on the LDDMM landmark manifold $(\mathbb{R}^{k\times d}, g_m)$. In the remainder of this section we give a brief description of the Riemannian metric $g_m$ and the induced distance metric $d_{g_m}$. For more details, see e.g. Chapter 4 of [1].

A starting point for LDDMM is to consider the space of *diffeomorphisms* on $\mathbb{R}^d$, i.e. smooth maps from $\mathbb{R}^d$ to itself, and a Riemannian metric on this space. This metric on diffeomorphisms induces a Riemannian metric $g_m$ on $\mathbb{R}^{d\times k}$, and thereby also a geodesic distance metric $d_{g_m}$ on $\mathbb{R}^{d\times k}$. The distance between two landmark matrices $x, y \in \mathbb{R}^{d\times k}$, defined in this way, can be interpreted as the least 'energy' needed to deform $\mathbb{R}^d$ in such a way that each landmark in $x$ coincides with the corresponding landmark in $y$.

The LDDMM Riemannian metric can be expressed relatively explicitly. More specifically, we can explicitly express the *cometric*, $g_m^\star$, whose matrix representation is the inverse of the metric matrix. The cometric at $x \in \mathbb{R}^{d\times k}$ is a block matrix

$$g_m^\star(x) = [(K_x)_{i,j}]_{i,j=1..k}\,, \qquad \text{for} \quad (K_x)_{ij} = \kappa(q_i, q_j) I_d \in \mathbb{R}^{d\times d}$$

where $x = [q_1, \ldots, q_k]$ is a matrix consisting of $k$ landmarks $q_i \in \mathbb{R}^d$, $I_d$ is the $d$ by $d$ identity matrix and $\kappa$ is a *kernel function*. See [4] for a derivation. A standard choice is a Gaussian kernel $\kappa(q_i, q_j) := \exp^{-\|q_i - q_j\|^2/2\sigma^2} \in \mathbb{R}$ with parameters $\beta, \sigma > 0$ (see [1], [2], [5]). The kernel determines how much energy it takes to move nearby landmarks in an *un*correlated way; for larger $\sigma$'s, nearby points $x, y \in \mathbb{R}^d$ tend to be mapped to nearby points $\phi(x), \phi(y) \in \mathbb{R}^d$ by low-energy deformations $\phi$. The $\sigma$ parameter adds modelling flexibility, but also implies that there is no canonical metric on the LDDMM landmark manifold, as there is on Kendall's shape space. Thus, $\sigma$ is a hyperparameter.

For the LDDMM metric, there are no closed-form expressions for any of the Riemannian operations that we need for Algorithm 1. We therefore rely on methods for numerical integration of differential equations and optimization, leading to increased computational costs. The necessary Riemannian operations are implemented, based on automatic differentiation, in our Python library *jaxgeometry* (*https://github.com/ComputationalEvolutionaryMorphometry/jaxgeometry*). Since the cometric, but not the metric, can be expressed fairly explicitly, it is computationally more efficient to compute geodesics and the Log map via the Hamiltonian equations. This means that the tangent vectors mentioned in Algorithm 1 are computed as cotangent vectors and handled accordingly. Tangents are in isomorphic correspondence with cotangents via the Riemannian sharp map and its inverse, so these are simply two different representations of the same objects.

**Mapping between cotangent spaces** In Algorithm 1, the standardized independent contrast between nodes (shapes) $x_i, x_j \in \mathbb{R}^{d \cdot k}$ is computed as

$$s_{ij} = \frac{\mathrm{Log}_{x_i}(x_j)}{e_i + e_j},$$

where $e_i, e_j$ are the lengths of the branches leading to their common parent node and Log is the Riemannian logarithm. As mentioned in the previous subsection, we will consider $s_{ij}$ to be a cotangent (momentum) vector in the cotangent space $T^\star_{x_i}\mathbb{R}^{d\times k}$ at $x_i$. In order to compute the phylogenetic covariance matrix estimate $\Sigma$ (Eq. (0.1)), we need to take outer products of these standardized contrasts. This is only possible if they all belong to the same cotangent space, thus we need to map them to a common space - done in step *9* of the algorithm. This mapping can introduce distortions (see [6] Section 2.3.1) if the cotangent spaces are based at points far apart. Since the root estimate $\hat{r}$ is a type of mean value, it is natural to map the standardized contrasts to the cotangent space at this point. Thus $s_{ij} \in T^\star_{x_i}\mathbb{R}^{d\times k}$ is mapped to $\tilde{s}_{ij} \in T^\star_{\hat{r}}\mathbb{R}^{d\times k}$.

On the LDDMM landmark manifold, a linear map between cotangent spaces can be constructed as follows. Let $t \mapsto \phi_t, t \in [0,1]$, be the optimal path of diffeomorphisms which takes landmark $x^m \in \mathbb{R}^d$ of shape $x = [x^1, ..., x^k] \in \mathbb{R}^{dk}$ to landmark $y^m \in \mathbb{R}^d$ of shape $y = [y^1, ..., y^k] \in \mathbb{R}^{dk}$ , that is, $\phi_1(x^m) = y^m$ for each $m = 1..k$. Let $D\phi_1(x^m) \in \mathbb{R}^{d\times d}$ be the Jacobian matrix of $\phi_1$ evaluated at $x^m$. Let $A_{y\to x} \in \mathbb{R}^{dk\times dk}$ be the block-diagonal matrix whose $m$'th diagonal element is

$$[A_{y\to x}]_{m:(m+d),m:(m+d)} = (D\phi_1(x^m))^T , \quad m = 1\ldots k \tag{1}$$

Then $A_{y\to x} : T^\star_y\mathbb{R}^{dk} \to T^\star_x\mathbb{R}^{dk}$ is a linear map taking a momentum vector from the cotangent space at $y$ to the cotangent space at $x$[2][7]. Notice that this map depends only on the diffeomorphism at time 1, not on the geodesic path of diffeomorphisms from time 0 to 1. Thus, for any other diffeomorphism $\psi$ satisfying $\psi = \phi_1$ around a neighbourhood of each landmark $x^m, m \in \{1, \ldots, k\}$, the Jacobian and therefore the cotangent maps coincide. Regarding numerics, the Jacobians $D\phi_t(x^m)$ for each time $t \in [0,1]$ can be computed by solving an ODE (see [2]), which is significantly more numerically stable than computing Jacobians by, e.g., finite differences.

For Algorithm 1, we compute $\tilde{s}_{ij}$ by mapping the cotangent vector $s_{ij}$ from cotangent space $T^\star_{x_i}\mathbb{R}^{dk}$ to $T^\star_{\hat{r}}\mathbb{R}^{dk}$. Let $n_1, n_2, ..., n_{l-1}, n_l$ be the sequence of intermediate inner nodes constituting the unique shortest path between $n_1 = \hat{r}$ and $n_l = x_i$. Let $\phi_1^2, ..., \phi_1^l$ be the sequence of optimal diffeomorphisms, where $\phi_1^j$, $j \in \{2, .., l\}$ takes each landmark $m = 1..k$ of parent node $n_{j-1}$ to the corresponding landmark of child node $n_j$, i.e. $\phi_1^j(n_{j-1}^m) = n_j^m$. Then the composition of these diffeomorphisms takes the root to the node $x_i$, i.e.

$$\phi_1^l \circ \cdots \circ \phi_1^2(\hat{r}) = x_i. \tag{2}$$

By composing the corresponding Jacobians we can map the independent contrast $s_{ij}$ to the cotangent space at the root;

$$\tilde{s}_{ij} = A_{n_2 \to n_1} \circ \cdots \circ A_{n_l \to n_{l-1}} \cdot s_{ij} = A_{n_2 \to \hat{r}} \circ \cdots \circ A_{x_i \to n_{l-1}} \cdot s_{ij}.$$

### 2 Comments on the diffeomorphic independent contrasts algorithm

**Computations** of differential geometric quantities such as *Exp* and *Log* are done using our Python library *Jaxgeometry* (https://github.com/ComputationalEvolutionaryMorphometry/jaxgeometry). As described in Section 1 of this appendix, it is computationally more efficient to compute geodesics (*Exp* and *Log*) from the Hamiltonian equations. Therefore, in our numerical implementation, the tangent vectors in Algorithm 1 are computed as cotangent vectors, and the tangent spaces are cotangent spaces.

**Diffeomorphic independent contrasts (Algorithm 1) works for other shape representations than LDDMM** The algorithm works for leaf-nodes taking values on an arbitrary finite dimensional Riemannian manifold - e.g. various shape spaces. An important example is Kendall's shape space, which is a backbone of geometric morphometrics [8]. However, step 9. of the algorithm might need to be adapted to the manifold in question. If there's is a Lie group structure, as in the LDDMM case, one has available a path-independent map between tangent spaces via the differential of the group action. In the absence of such a path-independent map, parallel transport along geodesics can be used (for more information on this approach, see ([6], Section 2).

### 3 Stochastic model for simulating shapes

The stochastic model we use for simulating shapes was introduced in [9], [10] and, with more mathematical details as well as biological interpretations, in [11]. In this section we give a brief presentation of the process, for more details see [11].

We model the random evolution of landmark shapes by means of a certain stochastic process. The process can be described at different levels of mathematical abstraction, in terms of either finite or infinite dimensional objects. The finite dimensional description is used for practical implementations, while the infinite dimensional description adds certain theoretical guarantees. In this section, we focus on the finite dimensional description.

We model a shape as a set of $k$ points (landmarks) $x_1, \dots, x_k$ in a domain $D \subset \mathbb{R}^d$, for $d = 2$ or 3. In evolutionary biology, the Brownian motion is used as a very basic model for random evolution of some trait of interest. However, if we let each landmark evolve as independent Brownian motions, the collection of points will likely evolve into something that cannot represent biological shapes. We need the motion of different landmarks to be correlated in a way that depends on their distance; nearby landmarks should be more likely to move in the same direction compared to landmarks that are further apart. Below, we describe a stochastic process that achieves this.

Let $p_1, \dots, p_J$ be $J$ points on a grid in $D \subset \mathbb{R}^d$. To each point $p_j, j = 1 \dots J$, we associate a kernel function $k_{p_j} : D \to \mathbb{R}$ centered at $p_j$. In this work, we choose a scaled Gaussian density $k_{p_j}(x) = \alpha \cdot e^{-\frac{\|p_j - x\|^2}{2\sigma^2}}$, where $\alpha$ is the *amplitude* and $\sigma$ the standard deviation, or *width*, of the kernel. The value $k_{p_j}(x)$ is largest for $x = p_j$ and decreases with the distance between $x$ and $p_i$. We use this to construct a system of stochastic differential equations (SDE's) for landmarks $X^i \in D$, $i = 1 \dots k$, which associates a Brownian motion to each grid point $p_j$ and weighs it by the kernel value $k_{p_j}(X^i)$:

$$\begin{aligned} dX_t^1 &= \sum_{j=1}^{J} k_{p_j}(X_t^1) I_d \circ dW_j, \\ &\vdots \\ dX_t^k &= \sum_{j=1}^{J} k_{p_j}(X_t^k) I_d \circ dW_j. \end{aligned} \tag{3}$$

Here $W_j$ is a $d$-dimensional standard Brownian motion and $I_d$ is the $d$-dimensional identity matrix. '$\circ$' denotes a Stratotonovich stochastic integral (which is equivalent to an Itô integral with a certain drift term). In the remainder of this section, we explain why solutions to this SDE yields the desired correlated motion of landmarks described above.

Firstly, note that the amplitude $a$ scales the kernel and thus the Brownian motions, and thereby it affects the speed of the process. Regarding the width parameter, $\sigma$, we first note that each landmark is driven by the same set of $J$ Brownian motions. This implies that, if two landmarks satisfy $k_{p_j}(X_t^1) \approx k_{p_j}(X_t^2)$ for all $j \in \{1, \dots, J\}$ then their trajectories will be similar around time $t$. Specifically, their difference satisfies the SDE

$$d(X_t^1 - X_t^2) = \sum_{j=1}^{J} \left( k_{p_j}(X_t^1) - k_{p_j}(X_t^2) \right) I_d \circ dW_j,$$

implying that if $k_{p_j}(X_t^1) \approx k_{p_j}(X_t^2)$ for each $j$ then the difference will be close to constant - i.e. the landmarks move in parallel. $k_{p_j}(X_t^1)$ is close to $k_{p_j}(X_t^2)$ for all $j$ if the landmarks $X_t^1$ and $X_t^2$ are close to each other *relative to the width* of the kernel function. The more narrow the kernel is, the closer landmarks need to be in order for their displacements to be parallel, and vice versa, a wide kernel implies that landmarks further apart will be displaced parallelly. See also [12] for an expression of the covariance between $X^1$ and $X^2$, which implies the same behavior. See Figure 1 for an illustration of the effect of different kernel widths. Each figure shows the position of 200 landmarks at time 1 (blue dots) after following the SDE (3) initialized in the circle configuration (black dots). The kernel width (standard deviation) was $\sigma = 0.1$ for the left figure, $\sigma = 0.3$ for the middle and $\sigma = 1$ for the right figure. The amplitude was $a = 1/20$ in all cases.

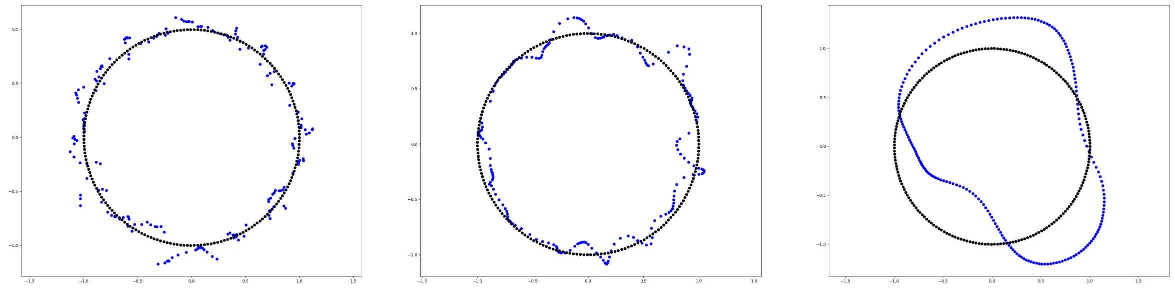

Figure 1: In each figure, the black dots show initial positions of 200 landmarks. The blue dots show landmark positions at time $t = 1$ after following SDE (3) with kernel parameters $a = 1/20$ and $\sigma = 0.1$ (left figure), $\sigma = 0.3$ (middle figure), $\sigma = 1$ (right figure).

The (3) in fact induces a stochastic path of diffeomorphisms on $D$. That is, the map from initial state to the state of a solution at time $t$ is a diffeomorphism. This implies that the model is consistent with the way shapes are treated in the LDDMM framework for shapes. For more details, see [... Sofias paper].

### 4 Landmark Methodology

The segmented images were used to manually place six anatomical landmarks with the function *digitize2D* from *Geomorph* ([13]) in R, to differentiate between morphological features such as hind-/fore-wings and thorax. Using *cv2.findContours* from *opencv* ([14]) in Python, the segmented image was used to extract the contour as points. These points were matched with the six anatomical landmarks, with the closest points in the outline classified as new anatomical landmarks. The index of the outline point was adjusted to initiate the sequence at the anatomical landmark between the left fore- and hind-wing. The outline was then anatomical into six regions between the anatomical landmarks. The region defining the head/antenna and thorax/tail points was removed, leaving four regions (two hindwings and two forewings), with a shared point at the inner angle between the hind- and forewing.

The total Euclidean distance across each region was accumulated point by point. $M$ equidistant landmarks were positioned on the outline using *numpy.linspace*, from 0 to the total distance, with M steps. The positions were translated into (x,y) coordinates by iterating over the region, finding the points before and after each distance position, and interpolating between these to reach the exact positions. A value of $M = 30$ landmarks was selected to provide a comprehensive representation of wing morphology and curvature while maintaining appropriate spacing for smaller wings.

Due to the absence of comprehensive databases for swallowtail and swordtail classifications across Papilionidae genera, a manual categorization system was implemented. Each specimen was classified based on tail morphology into three categories: un-tailed, tailed, or long tail. The dataset was individually aligned using general Procrustes alignment [15].

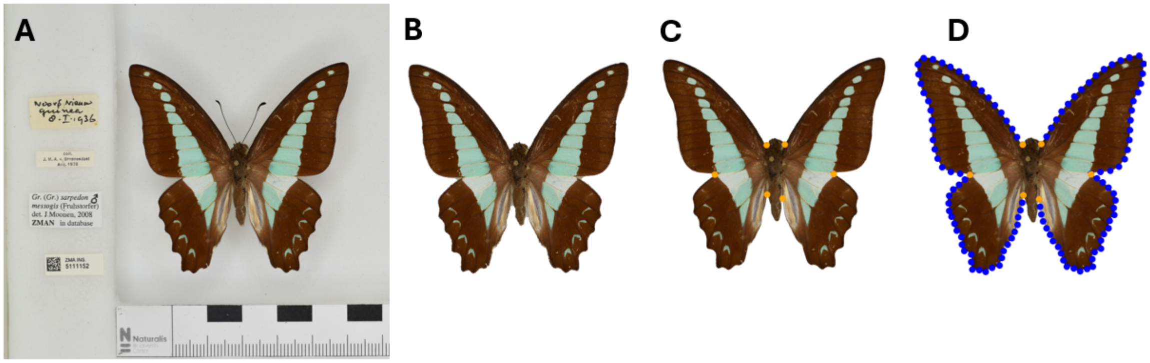


Figure 2: (A) The full image of a *Graphium Sarpedon*, which have been obtained from *Naturalis Biodiversity Center* through GBIF.org (B) The background are segmented out using *Segment Anything* in Python (C) Six manually placed anatomical landmarks (in orange) (D) 30 equidistance landmarks (in blue) are sampled along the outline between each of the manual placed landmarks. These are the points which are used for the subsequence analysis.

#### 5 Mean shapes

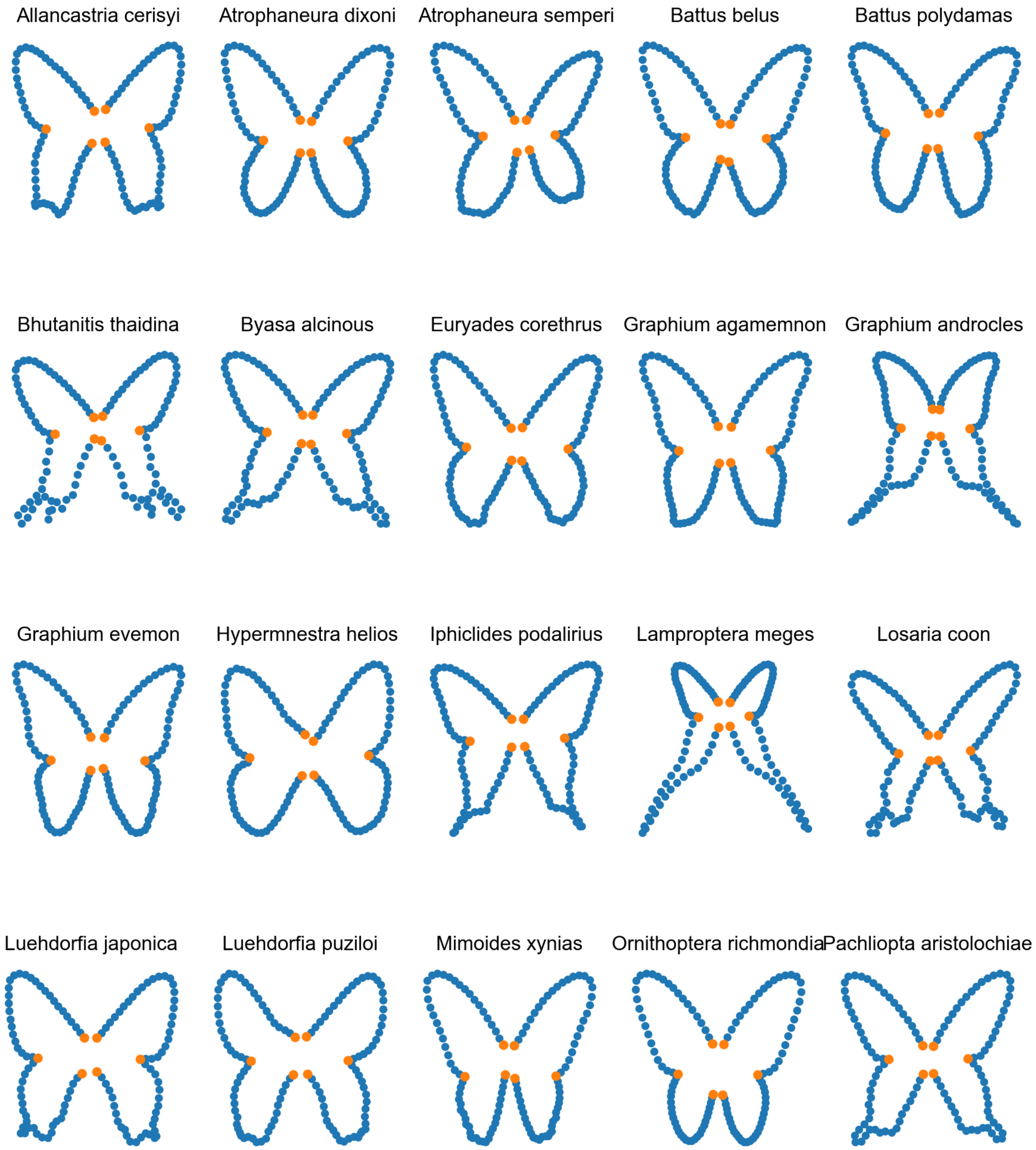

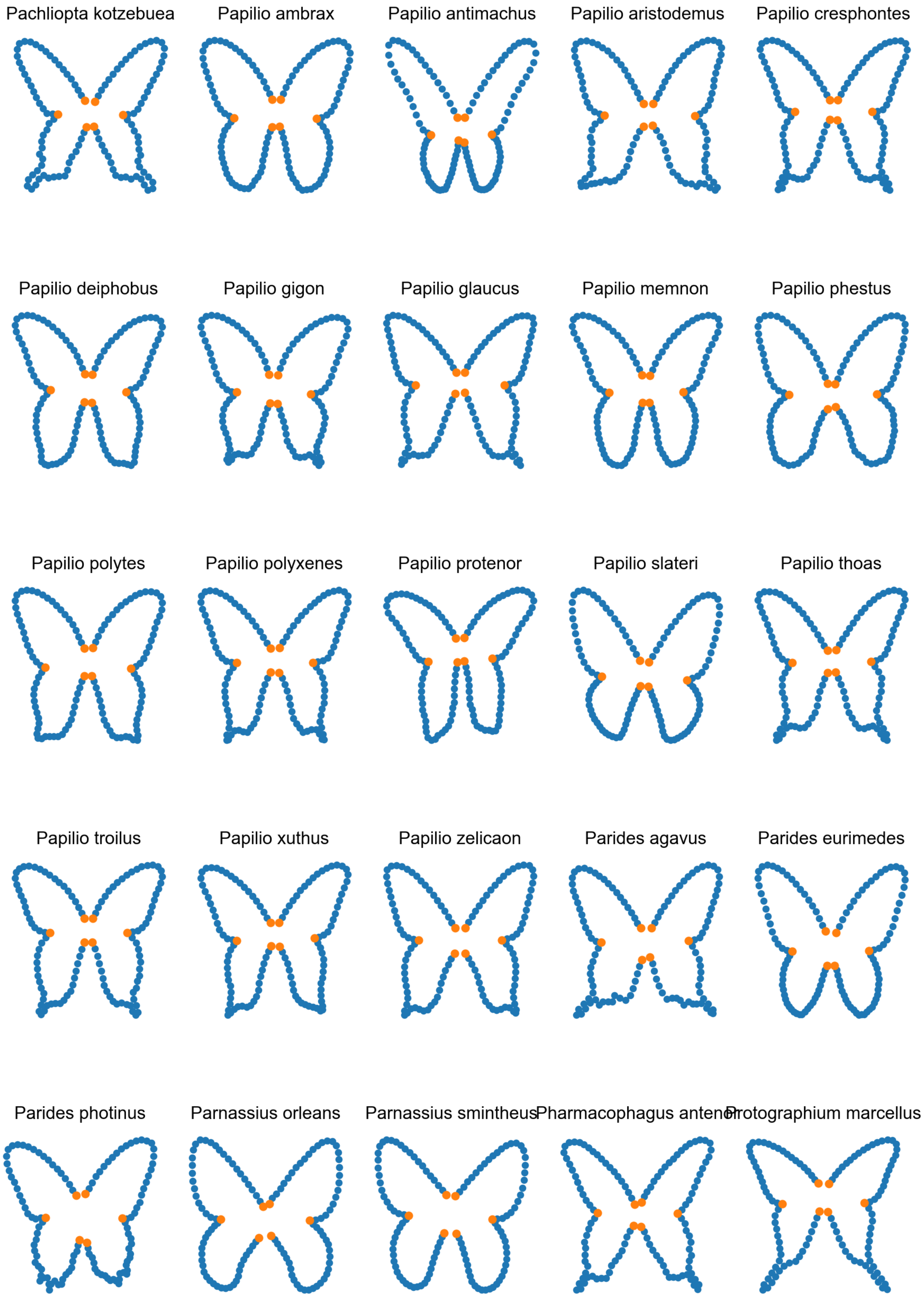
Pachliopta kotzebuea
Papilio ambrax
Papilio antimachus
Papilio aristodemus
Papilio cresphontes
Papilio deiphobus
Papilio gigon
Papilio glaucus
Papilio memnon
Papilio phestus
Papilio polytes
Papilio polyxenes
Papilio protenor
Papilio slateri
Papilio thoas
Papilio troilus
Papilio xuthus
Papilio zelicaon
Parides agavus
Parides eurimedes
Parides photinus
Parnassius orleans
Parnassius smintheus
Pharmacophagus anten
Protographium marcellus

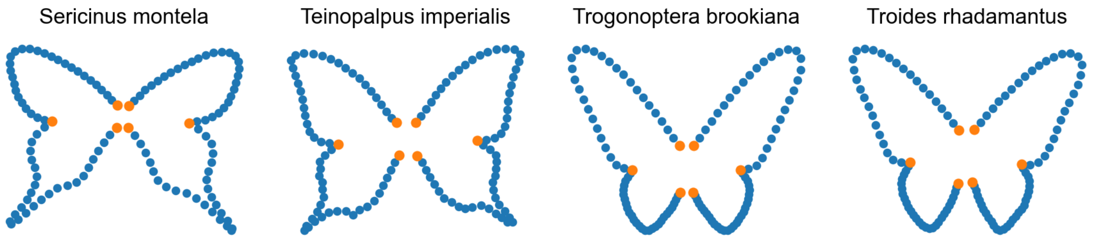
Sericinus montela
Teinopalpus imperialis
Trogonoptera brookiana
Troides rhadamantus

### 6 Origin of transformed images

| gbifID | license | publisher | catalogNumber | scientificName |
|---|---|---|---|---|
| 1321425831 | CC0_1_0 | National Museum of Natural History, Smithsonian Institution | USNMENT00781121 | Papilio belus Cramer, 1777 |
| 1436114194 | CC_BY_NC_4_0 | Museum of Comparative Zoology, Harvard University | 134885 | Sericinus montela Gray, 1852 |
| 863595061 | CC_BY_NC_4_0 | Museum of Comparative Zoology, Harvard University | 134830 | Allancastria cerisyi |
| 1438640363 | CC_BY_NC_4_0 | Museum of Comparative Zoology, Harvard University | 135072 | Teinopalpus imperialis Hope, 1843 |
| 1319561610 | CC0_1_0 | National Museum of Natural History, Smithsonian Institution | USNMENT00658629 | Euryades corethrus (Boisduval, 1836) |
| 4060704921 | CC0_1_0 | Naturalis Biodiversity Center | ZMA.INS.5149074 | Papilio gigon Felder & Felder, 1864 |
| 4406888457 | CC0_1_0 | Naturalis Biodiversity Center | ZMA.INS.5149573 | Papilio phestus Guérin-Méneville, 1830 |
| 3042580213 | CC_BY_NC_4_0 | Museum of Comparative Zoology, Harvard University | 211248 | Papilio antimachus Drury, 1782 |

### References


[1] Xavier Pennec, Stefan Sommer, and Tom Fletcher. *Riemannian geometric statistics in medical image analysis*. Academic Press, 2019.

[11] Sofia Stroustrup et al. "Stochastic phylogenetic models of shape". In: *In prep* ().

[12] Holm D.D. Arnaudon A. and S. Sommer. "Stochastic flows and shape bridges". In: *Statistics of Stochastic Differential Equations on Manifolds and Stratified Space* 48 (2021), pp. 18–21.

[13] Dean C Adams and Erik Otárola-Castillo. "geomorph: an R package for the collection and analysis of geometric morphometric shape data". In: *Methods in ecology and evolution* 4.4 (2013), pp. 393–399.

[14] Gary Bradski and Adrian Kaehler. *Learning OpenCV: Computer vision with the OpenCV library*. " O'Reilly Media, Inc.", 2008.

[15] John C Gower. "Generalized procrustes analysis". In: *Psychometrika* 40 (1975), pp. 33–51.
